# Proteome-wide 4-hydroxy-2-nonenal signature of oxidative stress in the marine invasive tunicate *Botryllus schlosseri*

**DOI:** 10.1101/2024.07.19.604351

**Authors:** Dietmar Kültz, Alison M. Gardell, Anthony DeTomaso, Greg Stoney, Baruch Rinkevich, Andy Qarri, Jens Hamar

**Affiliations:** University of California Davis, Department of Animal Sciences & Genome Center, Meyer Hall, One Shields Ave., Davis CA 95616, USA; University of Washington Tacoma, School of Interdisciplinary Arts and Sciences, 1900 Commerce St., Tacoma WA 98402, USA; University of California Santa Barbara, Department of Molecular, Cellular and Developmental Biology, Life Sciences Building 235, UCEN Rd, Goleta CA 93117, USA; Israel Oceanography & Limnological Research, National Institute of Oceanography, POB 9753, 3109701 Haifa, Israel; Helmholtz Zentrum München, Regenerative Biology and Medicine Institute, Munich, Germany

**Keywords:** Posttranslational modification, Lipid peroxidation, Environmental stress, Ecological proteomics, Invasive species, Colonial tunicates, Chordata

## Abstract

The colonial ascidian *Boytryllus schlosseri* is an invasive marine chordate that thrives under conditions of anthropogenic climate change. We show that the B. schlosseri expressed proteome contains unusually high levels of proteins that are adducted with 4-hydroxy-2-nonenal (HNE). HNE represents a prominent posttranslational modification resulting from oxidative stress. Although numerous studies have assessed oxidative stress in marine organisms HNE protein modification has not previously been determined in any marine species. LC/MS proteomics was used to identify 1052 HNE adducted proteins in B. schlosseri field and laboratory populations. Adducted amino acid residues were ascertained for 1849 modified sites, of which 1195 had a maximum amino acid localization score. Most HNE modifications were at less reactive lysines (rather than more reactive cysteines). HNE prevelance on most sites was high. These observations suggest that B. schlosseri experiences and tolerates high intracellular reactive oxygen species levels, resulting in substantial lipid peroxidation. HNE adducted B. schlosseri proteins show enrichment in mitochondrial, proteostasis, and cytoskeletal functions. Based on these results we propose that redox signaling contributes to regulating energy metabolism, the blastogenic cycle, oxidative burst defenses, and cytoskeleton dynamics during B. schlosseri development and physiology. A DIA assay library was constructed to quantify HNE adduction at 72 sites across 60 proteins that represent a holistic network of functionally discernable oxidative stress bioindicators. We conclude that the vast amount of HNE protein adduction in this circumpolar tunicate is indicative of high oxidative stress tolerance contributing to its range expansion into diverse environments.

**NEW & NOTEWORTHY:** Oxidative stress results from environmental challenges that increase in frequency and severity during the Anthropocene. Oxygen radical attack causes lipid peroxidation leading to HNE production. Proteome-wide HNE adduction is highly prevalent in *Botryllus schlosseri*, a widely distributed, highly invasive, and economically important biofouling ascidian and the first marine species to be analyzed for proteome HNE modification. HNE adduction of specific proteins physiologically sequesters reactive oxygen species, which enhances fitness and resilience during environmental change.

## INTRODUCTION

Diverse environmental challenges that cause oxidative stress increase in frequency and severity during the Anthropocene, including in the oceans (1). Oxidative stress is a universal secondary consequence of many other types of stresses in most organisms (2). Therefore, quantifying and interpreting functional consequences of oxidative stress, especially in marine organisms, remains a challenging but critical task for understanding the consequences and future trajectory of global anthropogenically induced environmental changes (3).

Tunicates are a chordate subphylum that represents the closest paraphyletic relative of vertebrates (4). Because of their close phylogenetic relationship to vertebrates, tunicates are models for studying effects of environmental stress on marine organisms (5, 6) and how the immune system has evolved in chordates to give rise to adaptive immunity in vertebrates (7). Moreover, despite being closely related to vertebrates, some colonial tunicates, including the golden star tunicate (*Botryllus schlosseri*), undergo whole-body regeneration as part of their asexual reproductive cycle (8). This astonishing ability makes them unique models for biomedical and bioengineering studies that are geared towards limb regeneration (9).

Besides their role as models of environmental and biomedical research, marine tunicates also represent major ecological and economic problems as invasive species. *B. schlosseri* is an invasive species that has colonized most temperate and subtropical oceans of the world and has recently also been reported from tropical and subarctic seas (10). It forms colonies comprised of flower-shaped systems that share a common tunic (Supplementary figure S1). Furthermore, tunicates are major biofouling organisms that have large economic impacts on many maritime industries (11). Biofouling by marine ascidians also represents a significant challenge for aquaculture, including the induction of oxidative stress in cultured mussels (12). A major challenge regarding the development of antifouling agents is to maximize their selective toxicity for biofouling target species while minimizing detrimental effects of beneficial non-target species, e.g. species used for aquaculture.

Biocidal effects of antifouling agents have been tested by assessing oxidative stress responses of marine invertebrates and fishes (13–15). For instance, exposure of aquaculture catfish (*Clarias gariepinus*) to antifouling chemicals increased the blood cell concentration of lipid peroxidation products, which are precursors of 4-hydroxy-2-nonenal (HNE) (16). Within tunicates, exposure to the antifouling agent tributyltin inhibited cytoskeletal dynamics important for cell division in *Styela plicata* (17) and decreased levels of glutathione (reduced form) within hemocytes of *B. schlosseri* (18). Oxidative stress is a universal stress triggered secondarily by many other types of environmental stress (19). Oxidative stress is omnipresent in marine environments and changing as a result of anthropogenic activities (3). Therefore, quantitation of oxidative damage and antioxidant activity in surrogate species has been used as a bioindicator of environmental and organismal health (20–22). For example, *B. schlosseri* has been transplanted to different sites in the Lagoon of Venice to measure the environmental quality at those sites based on the mRNA abundances of the antioxidant enzymes glutathione synthase, gamma-glutamyl-cysteine ligase, glutathione peroxidases, catalase, and Cu/Zn superoxide dismutase (23). Monitoring oxidative stress responses in indicator species can inform management practices in aquaculture farms and industries that are significantly impacted by biofouling, e.g. maritime, naval, and offshore mining businesses (11, 24).

In this study we analyzed the level of HNE adduction of *B. schlosseri* proteins to gain insight into the relevance of this lipid peroxidation product as a novel biomarker of environmental stress in colonial tunicates. Tunicates and other marine ectotherms are particularly susceptible to lipid peroxidation because of their high content of polyunsaturated fatty acids (PUFAs) (25, 26). PUFAs are fragmented into electrophilic aldehydes by lipid peroxidation chain reactions (27, 28). The most abundant aldehyde produced by lipid peroxidation is 4-hydroxy-2-nonenal (HNE) with significant amounts of malondialdehyde also being produced (28, 29). These electrophilic aldehydes are highly reactive and have very short residence times before covalently binding to proteins and forming posttranslational HNE modifications at specific amino acid residues.

In general, HNE modified proteins are considered impaired in their tertiary structure and function, detrimental for cellular homeostasis, and enriched during pathological conditions such as neurodegenerative and autoimmune diseases (30). For example, HNE abundance on key enzymes of oxidative metabolism (heme oxygenase 1, ATP synthase alpha, and enolase) increases during Alzheimers disease (AD) (27). Even during preclinical stages of AD and other neurogenerative diseases such as amnestic mild cognitive impairment, HNE abundance increases on proteins that regulate glucose metabolism, mTOR activation, proteostasis, and protein phosphorylation (30–32). Protein adduction rather than depletion of reduced cellular antioxidant pools has been identified as the primary mechanism of HNE toxicity (28, 33). Importantly, HNE does not merely have a role in pathophysiological processes. Instead, ample evidence suggests that HNE, when present at low concentration, can promote cellular and organismal health by regulating critical physiological functions, including signal transduction, cell proliferation, cell differentiation, anti-cancer defense mechanisms, angiogenesis, cell adhesion, and programmed cell death (34–37).

The present study introduces a direct, systems level approach for quantifying oxidative stress responses involving HNE protein adduction in *B. schlosseri*, which can be readily adopted for other organisms. It represents the first report on HNE in any tunicate. The approach presented here measures the amount of HNE lipid peroxidation product that reacts with proteins in oxidatively stressed cells. Our approach permits distinction of the specific proteins and amino acid residues that are differentially susceptible to oxidative damage in diverse environmental contexts to facilitate interpretation of the functional and phenotypic consequences of context-dependent environmental change. This approach is used here to compare the extents of HNE PTMs on specific proteins and amino acid residues in *B. schlosseri* field populations with colonies grown *ex situ* for many generations under fixed laboratory culture conditions.

## MATERIALS AND METHODS

### Sample preparation and LC/MS acquisition

*B. schlosseri* samples were collected as described in late summer 2022 from two marinas located 1800 km apart, in Santa Barbara, CA (population W, n=12) and Des Moines, WA (population T, n=20) (38). A third population includes laboratory raised *B. schlosseri* colonies that were derived from gravid colonies of the Santa Barbara marina population and then propagated for one year in the laboratory using a seawater flow-through system at the University of California Santa Barbara as described (8). Samples from this laboratory raised stock (population C, n=12) were collected in parallel to those from the two field populations. Sample collection, homogenization, and protein extraction, reduction, and alkylation were performed as described (38). Proteins were then digested for 3h using 50:1 parts of sample protein to trypsin/Lys-C (Thermo), peptides purified with C18 cartridges (Thermo), and buffer exchanged to 0.1% formic acid in LC/MS water (Thermo) using a speedvac (Thermo). Complex peptide mixtures (200 ng per sample) were separated on a 3 – 33% acetonitrile 70 min gradient using a nanoElute2 / Impact II LC/MS system using columns and acquisition parameters as previously reported (38). BSA standards (12.5 fmol) were analyzed before and after the entire set of samples to serve as a quality control for consistent instrument performance throughout analyzing the entire sample set. The bovine reference proteome for analyzing BSA standards was the same as previously used (38).

### Identification of HNE modified proteins and site-specific localization of HNE

The analysis of data from data-dependent acquisition (DDA) was conducted using PEAKS Studio 10.6 using the *B. schlosseri* reference proteome, which included 50% decoys and common contaminants (38). Bruker raw data for nine reference samples (three different systems for each *B. schlosseri* population) were directly imported into PEAKS and subjected to *de novo* sequencing using the following parameters: 20 ppm parent mass error tolerance, 0.02 Da fragment mass error tolerance, trypsin as enzyme, Cys carbamidomethylation and Met oxidation as variable PTMs, with maximum three variable PTMs per peptide. Subsequently, all samples were searched against the *B. schlosseri* reference proteome (38) in two rounds. During the first round, the following parameters, along with those employed for *de novo* sequencing, were applied: monoisotopic mass search, zero max. missed cleavages, and specific (complete) digest mode. During the second round, all 312 PTMs integrated into PEAKS Studio 10.6 were allowed as variable PTMs, to prevent bias towards the identification of specific PTM types and to avoid obligating the search algorithm to match peptide masses to conform to a predetermined PTM. HNE was selected from the resulting PTM profile of the second-round search and results for all nine DDA samples were exported to an Excel file. These exported data included information for all detected HNE modified peptides, including assigned protein accession number, protein group cluster ID, amino acid position of the PTM in the protein sequence, and PTM localization confidence A score (Supplementary table 1).

### Generation of a DIA assay library for partly HNE modified peptides

PEAKS Studio 10.6 second round search results for the nine reference samples (three different systems for each *B. schlosseri* population) were exported as pepxml and mzxml files and imported into Skyline 23.1 (39) to generate a raw spectral library. This library was then filtered for HNE modified peptides and their corresponding unmodified peptide variants. All HNE modified peptides detected were evaluated for inclusion in a data-independent acquisition (DIA) assay library to enable accurate quantitation of PTM modification ratios on specific amino acid residues of each HNE modified protein. First, all peptides with less than the maximally possible A score of 1000 were excluded. Then, HNE modified peptides were excluded if no corresponding peptide with an unmodified residue at the same position as the HNE modification was detected, due to the inability to calculate a PTM ratio. Finally, peptides lacking at least four diagnostic transitions (unique MS/MS ions free of interferences) were excluded from the DIA assay library. After this filtering process 60 proteins containing 72 HNE modified peptides remained in the DIA assay library. The DIA raw data for all samples were then imported directly into Skyline and used in conjunction with this DIA assay library. The following full scan settings were used for Skyline import of DIA raw data: centroided product mass analyzer, results (0.5 margins) isolation scheme, 15 ppm mass accuracy, and only scans within 2 minutes of the MS/MS identification retention time collected.

### Quantitation of HNE modification ratios

Since each amino acid residue modified by HNE was represented by at least one modified and one unmodified peptide variant in the DIA assay library, it was possible to directly compute the modification (HNE PTM) ratio for each HNE-modified amino acid residue in all proteins. For this purpose, peak area intensity data were exported for all peptides from Skyline to Excel, and the corresponding modified to unmodified ratio was then calculated for each amino acid residue in Excel as described (40). Quantitative normalization between samples was deemed unnecessary as the ratios derive from quantitative values for both modified and unmodified peptides within the same sample. Thus, they are inherently normalized as both modified and unmodified peptides were detected in all samples. Mean PTM ratios for each *B. schlosseri* population and the standard error of the mean were calculated for each HNE modified amino acid residue in Excel. Statistical analysis for differences in PTM ratios between *B. schlosseri* populations were performed by first analyzing the data for homoscedasticity (homogeneity of variances) using an F test, which was followed by application of the appropriate two-tailed t-test (type 2 or 3). Modification ratios were considered significantly different between distinct *B. schlosseri* populations if p<0.05 (-log10 p>1.3) and the change of an HNE PTM ratio > 2-fold.

### String functional enrichment analysis of HNE modified proteins

A custom STRING (41) reference database, generated previously for *B. schlosseri,* was used for functional enrichment analysis (38). STRING network analyses were performed using the multiple protein search option. STRING clusters of proteins were considered enriched or depleted when the false discovery rate was less than 1% (FDR<0.01). Resulting STRING network clusters were visualized using default settings, except for displaying network edges as a single line. The thickness of the line indicates the confidence (strength) of the connection between protein nodes. Markov clustering (MCL) was performed with the inflation parameter set at 3. Nodes within the same MCL network cluster were connected by solid lines while nodes belonging to different network clusters were connected by dashed lines. Network clusters were color-coded to readily visualize protein nodes belonging to the same cluster.

## RESULTS

### HNE modification of the *B. schlosseri* proteome is extensive

Analysis of HNE modifications on a proteome-wide scale was performed by considering data for nine *B. schlosseri* samples analyzed by DDA (three samples from each population). This analysis revealed 1052 proteins containing at least one HNE-modified residue (1849 total modified residues, Supplementary table 1). Almost two-thirds (1195) of these HNE PTMs had an A score = 1000 (Figure 1a), which is the maximum possible A score representing the highest confidence for localization of a PTM to a specific amino acid residue (42). For the majority of these HNE modified residues (1185) no corresponding unmodified peptide was detected, indicating that the extent of HNE modification is massive in all three *B. schlosseri* populations analyzed. Nevertheless, slightly more than a third (664) of HNE modified residues were accompanied by peptides that were unmodified on the same residue, suggesting that different amino acids within specific proteins have unique propensities for being HNE modified. Moreover, HNE modification ratios can be quantified if both modified and unmodified versions of the peptide are robustly detectable above the instrument sensitivity limit.

**Figure 1:**
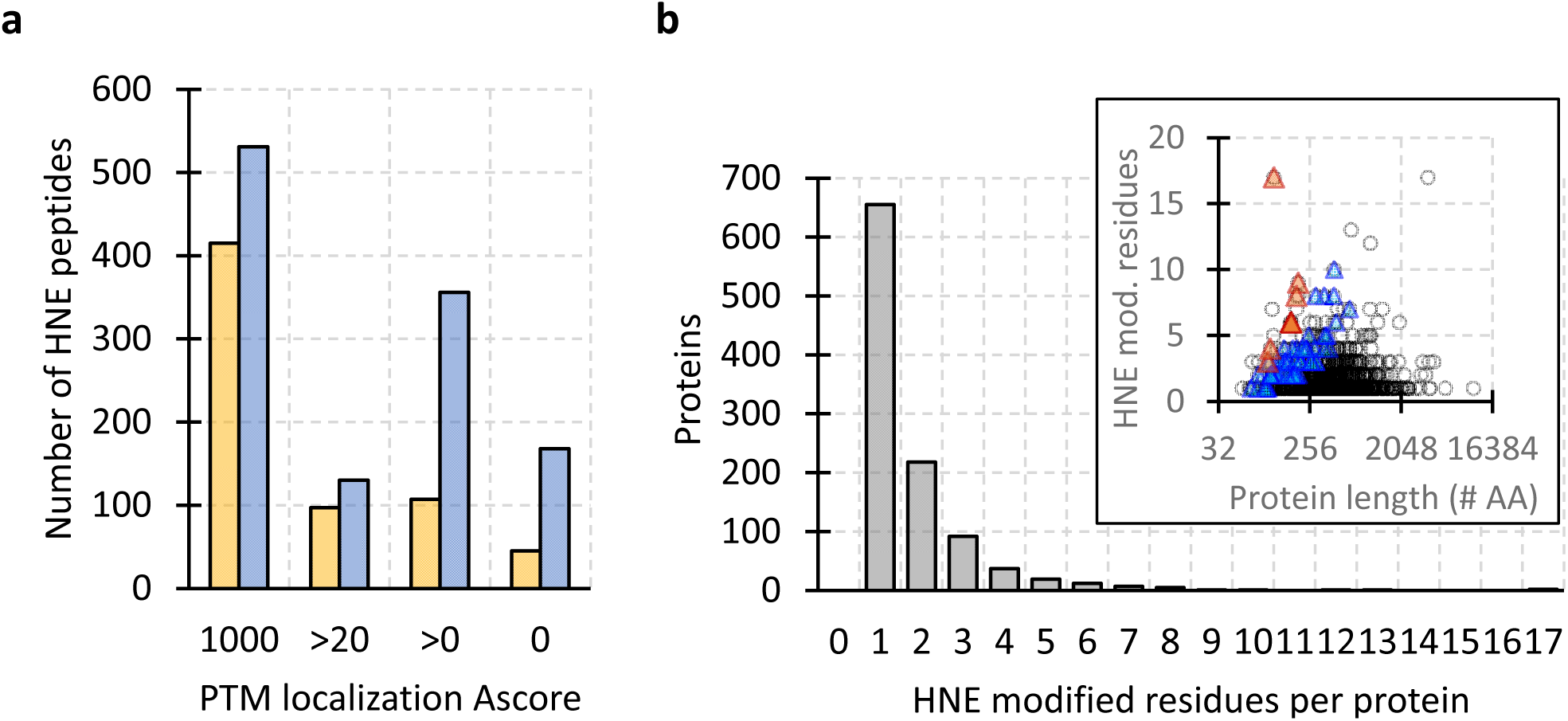
Number of HNE modified amino acid residues and proteins identified. A) Number of peptides with HNE modification depending on PTM localization A score. The max. A score of 1000 denotes the highest confidence of HNE assignment to a specific amino acid residue. Yellow bars = HNE modifications for which a corresponding peptide containing the unmodified amino acid residue was detected. Blue bars = HNE modifications for which no corresponding unmodified peptide was detected, i.e., all detected petide(s) containing the amino acid residue were HNE modified on that residue. B) Number of HNE modified amino acid residues per protein. The inset shows how the number of HNE modified residues corresponds to protein length (number of amino acid residues). Proteins with the greatest proportion of HNE modified residues having an A score of 1000 normalized by total number of amino acids (protein length) are depicted as orange (>3% HNE modified residues) and blue (>1% HNE modified residues) triangles.

Most proteins contained a single HNE modified residue, but some had more than 10 (Figure 1b). In general, larger proteins with more amino acids tended to have more HNE modified residues but there were notable exceptions of small proteins with many more than expected HNE modified residues (Figure 1b inset). The 14 proteins with the greatest enrichment in HNE modified residues (>3% of total amino acids in the protein sequence) included six mitochondrial (NADP-specific glutamate dehydrogenase, alanine-glyoxylate aminotransferase 2, fatty acyl-CoA hydrolase, medium chain-like enoyl-CoA hydratase, glutathione reductase, hydroxyacyl-coenzyme A dehydrogenase), two ribosomal (60S RPL9 and 40S RPS20), and five other (transcriptional activator protein Pur-beta, coactosin, cullin-associated NEDD8-dissociated protein 1, lactadherin, uncharacterized LOC120343960, unknown) proteins. An additional 120 proteins had an HNE enrichment >1% of total amino acids in the protein sequence and functionally reflected the set of 14 proteins with the greatest (>3%) HNE enrichment (mostly mitochondrial and ribosomal proteins, Supplementary table 2).

### Most HNE modifications are localized on lysine residues

More than half of the HNE modifications detected were on lysine followed by arginine, cystine, and histidine. This trend was even more striking for HNE PTMs with a maximum A score of 1000 (Figure 2a). The localization confidence of an HNE PTM to a particular amino acid residue (and corresponding A score) are based on a specific mass shift of 156.12 atomic mass units (amu) that is added to fragment (y and b) ions of the HNE modified peptide depending on the position of the HNE PTM in the peptide sequence (Figure 2b). In the example shown in Figure 2b this mass shift of 156.12 amu is evident for all y ions but not for b ions indicating that the HNE modification is present on the C-terminal lysine residue of the peptide. When only considering HNE PTMs with the maximal A score of 1000 then 507 Lys (53.6%), 389 Arg (41.1%), 32 Cys (3.4%), and 18 His (1.9%) residues are HNE modified in the *B. schlosseri* proteome of the three populations analyzed in this study.

**Figure 2:**
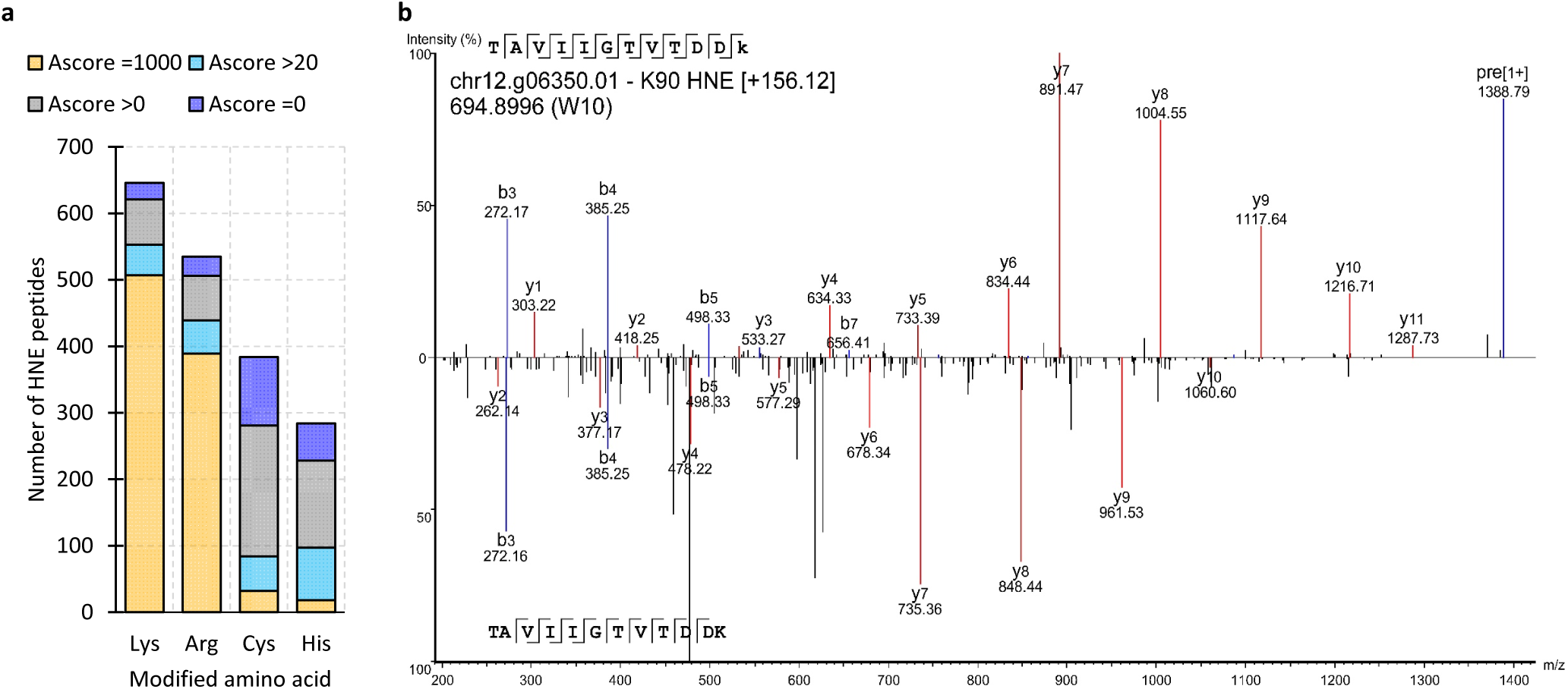
Localization of HNE modification on specific amino acid residues. A) Numbers of HNE modified Lys, Arg, Cys, and His residues depending on PTM localization A score (color-coded). B) Example MS/MS spectra for peptide TAVIIGTVTDDK containing C-terminal Lys90 of 60S ribosomal protein L18-B-like (chr12.g06350.01). The HNE modified MS/MS spectrum is shown above and the corresponding unmodified spectrum below the x axis. Unambiguous PTM localization to Lys90 is supported by a consistent mass shift of 156.12 amu in the entire y ion series (red lines) but no mass shift in the b ion series (blue lines).

### Functional enrichment of partly but not fully HNE modified proteins

A STRING functional enrichment analysis (41) was conducted to assess whether the fully HNE adducted proteins (where only the modified peptide was detected) exhibit similar functions compared to partly HNE-modified proteins (where both modified and unmodified peptides were detected). The proteins containing both fully and partly HNE modified resides (11.9% of all proteins with HNE PTMs having an A score of 1000, Figure 3a) were excluded from this analysis. The remaining 236 partly HNE modified proteins were strongly enriched for 13 functional STRING clusters. These clusters are postsynaptic cytoskeleton organization and positive regulation of actin filament polymerization (CL:3186 and CL:3187), actin cytoskeleton organization, and 7SK snRNA binding (CL:3184), Cytosolic ribosome (CL:1368, CL:1371, CL:1375 and CL:1380), structural constituent of ribosome and translation elongation factor activity (CL:1360), tricarboxylic acid cycle, and citrate metabolic process (CL:4849), mixed group, including muscle contraction, and intermediate filament organization (CL:6911), mixed group, including urea cycle, and 3 iron, 4 sulfur cluster binding (CL:4853), translation (CL:1358) and positive regulation of actin filament polymerization, and structural constituent of postsynaptic actin cytoskeleton (CL:3189) (Figure 3b). Actin cytoskeleton, ribosomal, and mitochondrial related functions are most evident in these clusters. The latter two of these three functional categories were also the most prominent functions for the set of proteins containing the most HNE modified residues per total number of amino acids (see above). Surprisingly, no significant functional enrichment was detectable for the set of 340 fully HNE modified proteins.

**Figure 3:**
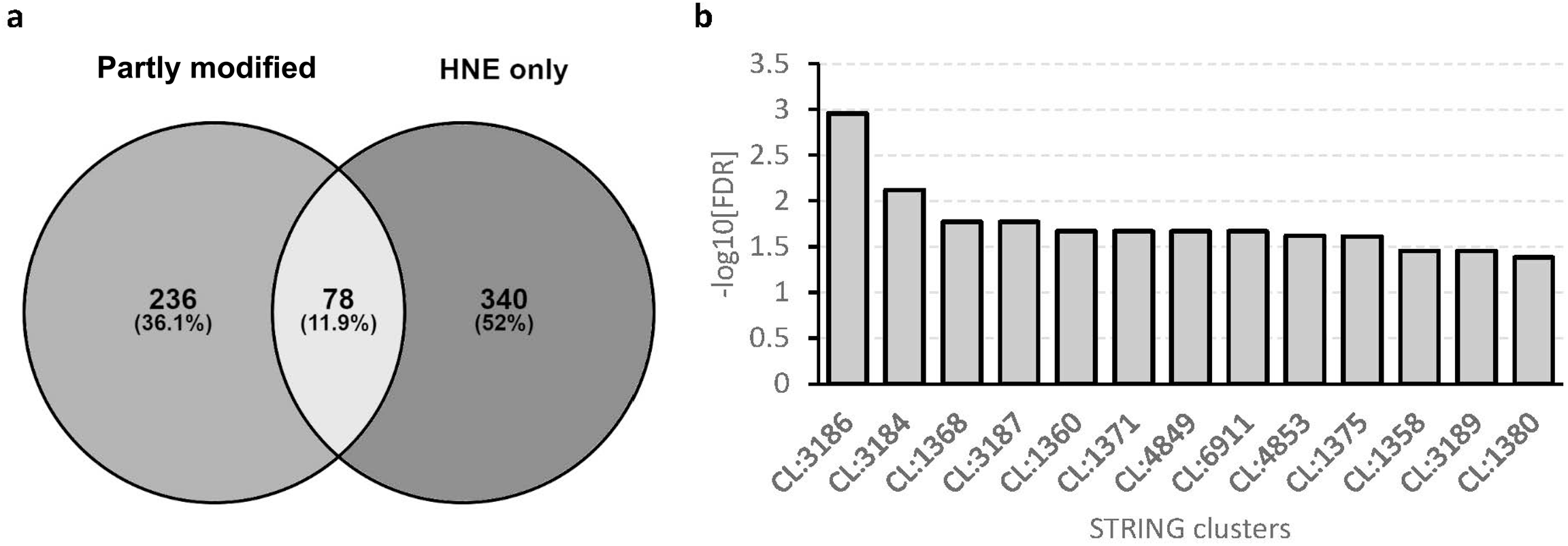
Functional enrichment analysis of HNE modified proteins with a PTM localization A score of 1000. A) Numbers of proteins with only partly HNE modified residues (left), proteins with two or more HNE modified residues, at least one of which is partly and another fully modified (center), and proteins with only fully HNE modified residues. B) Functional STRING clusters that are significantly enriched (FDR<0.01) for the set of 236 proteins that contain only partly HNE residues. The STRING cluster accession numbers are explained in the text.

### Proteins with quantifiable HNE modification ratios represent pertinent functions

Sixty proteins containing 72 partly modified HNE PTM residues for which modified and unmodified peptides were detected well above the sensitivity limit met all QC criteria for inclusion in a quantitative DIA assay library that included: an A score of 1000, at least four diagnostic transitions for each the modified and unmodified peptide variant, different retention times for modified and unmodified peptide variants, lack of ion interferences, and detectable quantities in all nine training samples. The same three samples that were used for the initial DDA runs for each population were used as training samples. The great majority of proteins included in the DIA assay library (n=50) contained a single HNE PTM site, nine proteins contained two HNE PTM sites, and one protein contained three HNE PTM sites. The proteins included in the DIA assay library reflect the dominance of Lys-HNE (52 residues) but also include 19 Arg-HNE, and 2 His-HNE residues. No Cys-HNE residues were included in this DIA assay library.

STRING MCL cluster analysis of the 60 proteins included in the DIA assay library revealed a network consisting of eight clusters (Figure 4a). Actin cytoskeleton, ribosomal, and mitochondrial related functions were again most evident in these clusters, confirming that the 60 proteins included in the DIA assay library are functionally representative of the majority of other HNE PTMs identified. The largest cluster contains 14 proteins involved in pre-mitochondrial carbohydrate and energy metabolism. The second largest cluster contains 10 proteins associated with intermediate filaments. The third largest cluster contains 9 proteins involved in ribosomal translation. The average HNE PTM ratios across all samples analyzed in this study were not cluster-specific but varied as much within as they did between clusters (Figure 4b). They ranged from less than 10% to over 80%. Fifteen proteins did not cluster together with any protein in the HNE protein network and eight proteins were not included in the network because of missing STRING annotations. The average HNE modification ratios of these proteins across all samples also varied between less than 10% to over 80% (Figure 4c).

**Figure 4:**
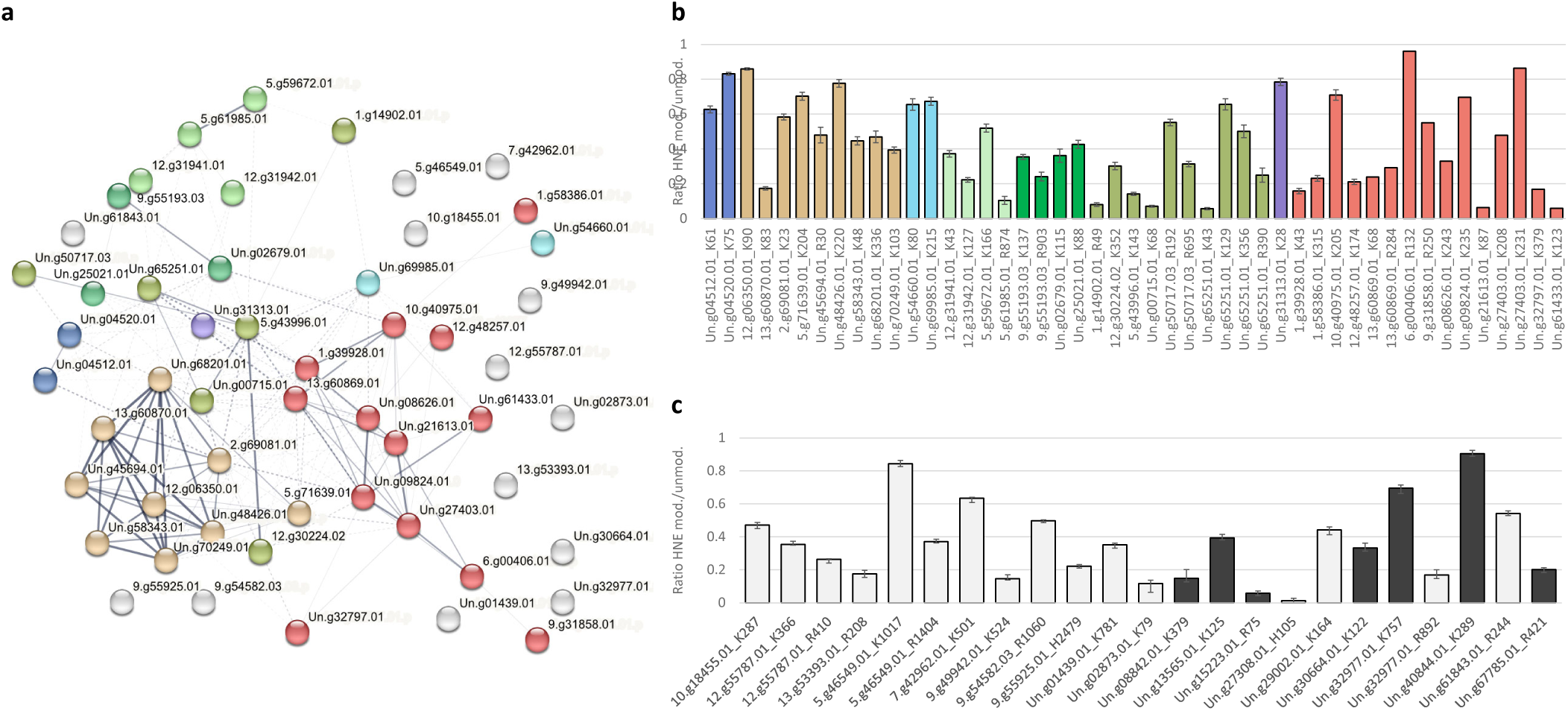
STRING analysis and PTM ratios of all HNE modified proteins with an A score of 1000 that met all criteria for inclusion in the quantitative DIA assay library. A) STRING network depicting functional relationships (edges) of HNE modified proteins (nodes). Functional STRING clusters are color-coded as follows: Red = Fructose/ mannose metabolism and pentose phosphate pathway; Beige = Translation at cytosolic ribosome; Olive green = Intermediate filament; Light green = Triglyceride metabolism and xenobiotic transport; Dark green = Actin filament depolymerization and growth factor activity; Blue = Regulation of actin filament polymerization by Rho GTPase signaling; Cyan = tubulin polymerization activity and immunoproteasome (no sign. enrichment); Purple = Glycogen synthesis (no sign. enrichment); B) Average HNE PTM ratios across all samples (mean ± SEM) for all residues of proteins that are colored and included in functional clusters depicted in panel A. Note that several of these proteins were HNE modified on more than one residue. C) Average HNE PTM ratios across all samples (mean ± SEM) for all residues of proteins not clustered with other proteins in panel A (white) or not included in the network in panel A because of missing STRING annotation (black).

### HNE modification ratios differ between laboratory reared and field populations

The effect of *ex situ* maintenance of *B. schlosseri* for over a year that included over 50 consecutive asexual generations of zooids on HNE modification ratios was assessed by comparing these ratios for each protein and residue represented in the DIA assay library between all three populations. Across all quantified HNE sites, the laboratory reared population C had 30 - 40% higher HNE modification ratios than both field populations. This trend was consistent for most HNE sites (Figure 5). In contrast, the overall difference between the two distant field sites (W and T) was less than 20% (Supplementary table 3). Nine HNE PTMs had significantly higher and three significantly lower ratios in the laboratory reared animals (C) than the corresponding parent population from the field (W) (Figure 5a). Each of these variable HNE PTMs was located on a separate protein. Significantly higher HNE modification ratios in C vs. W were found in WD repeat-containing protein 1 (chrUn.g02679.01), glial fibrillary acidic protein (chrUn.g65251.01), mitochondrial enoyl-CoA hydratase (chrUn.g21613.01), 60S ribosomal protein L3 (chrUn.g29002.01), armadillo repeat protein deleted in velo-cardio-facial syndrome (chr5.g61985.01), tubulin beta (chrUn.g08842.01) and three uncharacterized proteins (chr12.g55787.01, chrUn.g32977.01, chr9.g49942.01). The three proteins with significantly lower HNE modification ratios in C vs. W were actin (chrUn.g27308.01), fructose bisphosphate aldolase C (chr13.g60869.01), and an uncharacterized protein (chr12.g31941.01) (Supplementary table 3).

**Figure 5:**
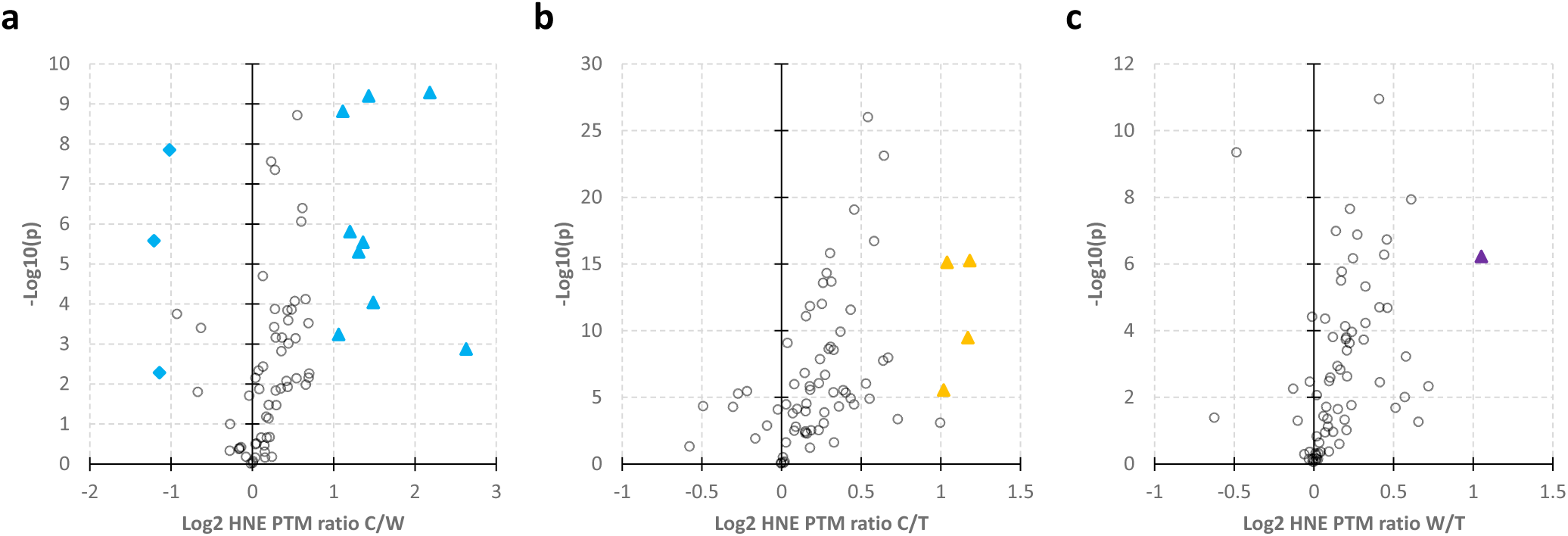
Volcano plots of HNE modification ratios compared between laboratory raised (C) and field (W and T) populations. A) Comparison of C versus W. Blue symbols represent significantly different (triangles = higher, diamonds = lower) HNE modification ratios. B) Comparison of C versus T. Yellow triangles represent significantly higher HNE modification ratios. C) Comparison of two field populations W versus T. A purple triangle represents a significantly higher HNE modification ratio.

Surprisingly, the difference between laboratory reared and field populations was less pronounced when comparing the C and T samples. Although many HNE modification ratios had highly significant p values, only four met the twofold change (2FC) cutoff and were significantly higher in C (Figure 5b). This result was unexpected as C was derived from W and T is located 1800 km apart from W. Three of these four HNE sites were significantly higher in both comparisons of laboratory versus field populations (C vs. W and C vs. T). These three HNE sites are present in glial fibrillary acidic protein (chrUn.g65251.01) and two uncharacterized proteins (chr12.g55787.01, chrUn.g32977.01). However, the HNE modification ratio on lamin-B1 (chr1.g14902.01) was not significantly increased in the C vs. W comparison (Supplementary table 1). Conversely, this HNE site on lamin B1 was the only residue with a significantly different PTM ratio when comparing two field populations that were collected 1800 km apart (Figure 5c).

## DISCUSSION

Only a limited number of studies have explored HNE modification of proteins or the regulation of free HNE in any aquatic invertebrates. Given the very high levels of HNE modified proteins in *B. schlosseri* it is noteworthy that HNE has not been documented in any tunicate prior to this study. In mussels (*Mytilus galloprovincialis*) HNE levels were found to be pollution- and sex-dependent (43). Exposure of the sessile marine polychaete *Galeolaria caespitosa* to dibutyl phthalate generated oxidative stress and increased HNE in sperm leading to alkylation of the sperm centrioles and disruption of cytoskeleton organization during early embryogenesis (44). In white-leg shrimp (*Litopenaeus vannamei*) mitochondrial uncoupling protein activity was stimulated by HNE (45). The above literature results are consistent with our finding of mitochondrial, energy metabolism enzymes, and cytoskeletal proteins being significantly overrepresented among partly HNE-modified *B. schlosseri* proteins. The set of fully HNE-modified proteins for which no unmodified peptide was detected also contains many proteins involved in these functions. No functions are overrepresented in this set possibly due to the presence of many novel, uncharacterized proteins among the fully HNE modified proteins that lack functional annotation and were not mapped to the STRING database.

The numerous fully HNE adducted proteins could potentially serve a protective function as antioxidant macromolecules that sequester high constitutive levels of lipid peroxides, arisen from the typical *B. schlosseri* lifestyle that incurs frequent exposure to elevated levels of oxidative stress. If that were the case, then natural selection may have yielded optimally functioning protein structures that include HNE modified residues as the preferred substrate during selection. This would mean that protein primary sequences and corresponding DNA coding sequences reflect optimal protein three-dimensional structures that include HNE PTMs at specific residues. Following this scenario, HNE modified proteins should function as well or even better than their unmodified counterparts, a hypothesis yet to be tested. This hypothesis contrasts with the typically detrimental consequences of HNE protein adduction leading to metabolic dysfunction in other species that are not routinely exposed to high levels of oxidative stress (46). Moreover, in such a scenario detoxification could potentially occur after proteolytic degradation of the proteins and expulsion of HNE-adducted Lys from the cell by amino acid transporters (47).

### HNE amino acid specificity indicates high and chronic oxidative stress in *B. schlosseri*

Physiological and cytotoxic consequences of HNE modification depend on which amino acid residue is modified, how such modification structurally and functionally impacts the corresponding protein, and to what extent the affected protein is essential (48–51). Lysine PTMs alter reversible protein-protein interactions, which have been shown to cause activity changes of histones, cytoskeletal proteins, and enzymes that regulate energy metabolism including mitochondrial enzymes (52, 53).

According to Pearson’s hard and soft acids and bases (HSAB) theory (54), HNE is considered a soft electrophile that primarily forms 1,4-Michael adducts with cysteine sulfhydryl thiolate groups as soft nucleophilic targets. Harder nucleophilic nitrogen groups of Lys, Arg, or His are thought to be three orders of magnitude less reactive with HNE than Cys (55). Consequently, it has been hypothesized that the cytotoxic effects of NHE and other aldehydes are primarily due to formation of Michael adducts with sulfhydryl thiolate groups of Cys in critical proteins (55). For mammals, these theoretical considerations have been supported by multiple proteomics studies (56–58).

Conversely, several other proteomics studies on mammals have shown that Lys is a preferred target of HNE adduction (59–62). HNE reacts with primary amines on Lys to form a Schiff base (63). It has been proposed that HNE modification of Cys may indicate low toxicity while HNE modification of Lys may indicate chronically high levels of oxidative stress (55). On this basis, our data are consistent with high levels of oxidative stress in all three *B. schlosseri* populations. However, our results also show that the extent of HNE protein adduction is significantly different between populations, which suggests that, although high in all populations, the level of oxidative stress varies between populations. Environmental parameters (temperature, pH, salinity, xenobiotic concentrations, etc.) vary significantly at each of the three source locations and likely contribute to the observed differences in HNE protein adduction.

The preferred amino acid reactivity of HNE may be species-specific, protein-specific, and/ or depend on the specific conditions inside the cell. pH greatly affects the rate of Cys reaction with type 2 alkenes (64). At a typical intracellular pH of 7.4 this reaction is 15 times slower than at pH 8.8 or the optimum that can be theoretically calculated (55). Moreover, Lys residues are, on average, more than twice as abundant (7%) in animal proteins than Cys residues (3%) (65). These considerations, along with our results, suggest that intracellular conditions in *B. schlosseri* favor adduction of HNE to Lys over Cys. The primary amine of Lys is also a target for various other PTMs that play a crucial role in cell signaling (acetylation, methylation, ubiquitination, etc.) (66). Thus, Lys HNE adduction may be cytotoxic by preventing other, reversible Lys PTMs that regulate cellular function and phenotype. Alternatively, HNE modification of proteins has been shown to represent a physiological mechanism that improves cell function and has an important redox signaling role in certain contexts (34, 37).

### Redox regulation of physiological functions in *B. schlosseri*

Oxidative stress and redox regulatory mechanisms play pivotal roles for various physiological functions in *B. schlosseri*. One example represents the outcomes of non-self recognition following direct tissue-to-tissue contacts between allogeneic incompatible partners that developed into what are called ‘points of rejection’ (POR), distinguished morphologically as dark-golden to brown cytotoxic lesions in the contact zone between adjacent partners (67). The formation of these PORs is triggered by the release of phenol oxidase and its polyphenol substrates from internal vacuoles of morula cells within 2h of recognizing chemical signals from a competitor colony (68). The cytotoxic effect of this enzyme can be prevented by the addition of phenol oxidase inhibitors, reducing agents, and antioxidant enzymes suggesting that reactive oxygen species (ROS) production leading to oxidative stress is the main cause of cytotoxicity during allogenic rejection (69).

Phenol oxidase is also involved in tunicate innate immune responses against parasites and pathogens. This enzyme and its substrates (polyphenols) are recruited to sites of infection from hemocytes that circulate throughout the vascular system and contribute to innate immunity. Phenol oxidase catalyzes the oxidation of polyphenols at sites of infection, which promotes production of inflammatory cytotoxic quinones, polymerization of melanin, and generation of ROS (7). Increased ROS levels, in turn, accelerate lipid peroxidation, which results in elevated HNE concentrations and an increase in HNE adducted proteins. It is therefore possible that the high levels of HNE adducted proteins in *B. schlosseri* are indicative of an active immune system that fends off pathogens and epiphytic organisms to prevent them from obscuring the tunic and attacking the zooid and vascular structures. Such microbial organisms may have an advantage relative to *B. schlosseri* under laboratory culture conditions because they have shorter generation times and can adapt to novel environments more rapidly.

Another example manifests redox regulation which is critical for the completion of the weekly *B. schlosseri* asexual life cycle, called blastogenesis. This asexual reproduction cycle includes a takeover phase during which adult zooids are resorbed through apoptosis and phagocytosis within 24 h, and are replaced by primary buds that develop to the new generation of functional zooids (70). This stage’s ‘life and death’ process is characterized by increased oxidative metabolism and concomitant increases in ROS concentration and antioxidant gene expression (71). During takeover apoptosis in zooids and resorption by circulating phagocytes is triggered via increased oxidative stress and prolonged respiratory burst (72), a physiological state of elevated reactive oxygen species production. Molecular chaperones, including proteins that are HNE modified, actively participate in the regulation of *B. schlosseri* blastogenesis and aging (73).

Cadherin has been implicated in the takeover process as it is upregulated during oxidative stress and other environmental stresses and its knockdown using siRNA during takeover disrupts morphogenesis and the blastogenic cycle (74). We have identified a cadherin homolog (AC # chrUn.g54100.01) among the HNE modified proteins in our study (Supplemental table 1). This cadherin is HNE adducted on Arg189. Moreover, three catenins that form a complex with cadherin that promotes cell adhesion are among the HNE adducted proteins identified in *B. schlosseri*. Beta-catenin (AC # chrUn.g33590.01.p) is HNE adducted on Lys10, beta1-catenin (AC # chr10.g39566.01.p) on Arg379, His430, and Cys459, and alpha2-catenin on Arg217 and Lys509 (Supplemental table 1). It would be interesting to compare the profiles of HNE adduction of cadherin and catenins at different developmental stages in future studies. Cadherin, catenins, and other HNE modified signaling and cytoskeletal proteins detected in *B. schlosseri* likely contribute to the regulation of programmed cell death, cytoskeleton dynamics, the takeover process, allogeneic incompatibility, and/or innate immunity in *B. schlosseri*.

### Enrichment of HNE modified mitochondrial proteins as indicators of oxidative stress

HNE adduction of proteins involved in energy metabolism, mitochondrial function, and cytoskeletal structure has been previously reported from other organisms with a suggested role in structural and functional destabilization of cells as a result of oxidative stress (29, 55). Increased ROS concentrations result from increased oxidative mitochondrial metabolism and activation of oxidase enzymes that are triggered by many types of environmental stress (2). Oxidative stress is a secondary type of stress that is an indirect consequence of many other types of environmental stress (19). It is therefore not surprising that HNE increases have been reported after exposure of red cusk-eels (*Genypterus chilensis*) to thermal stress (75) and in mussels (*Mytilus galloprovincialis*) exposed to pollution (43). During stress, mitochondrial energy metabolism increases to generate chemical energy equivalents (i.e., ATP) for fueling the cellular stress response. The increase in cellular respiration results in concomitant increases of mitochondrial ROS (2), causing subsequent increases in mitochondrial membrane lipid peroxidation (76, 77), which is consistent with the enrichment of HNE-adducted mitochondrial proteins observed in our study. Mitochondrial proteins are in close vicinity to free HNE released from mitochondrial membranes and may therefore be preferred targets of HNE adduct formation.

Free HNE and other free aldehydes generated by lipid peroxidation have much shorter half-lives than HNE adducted proteins, which are much more stable (29, 37). Therefore, measuring the levels of HNE adducted proteins and quantifying HNE modification ratios at specific amino acid residues of adducted proteins represents a more robust and reliable assay of oxidative stress than determination of free HNE. Sample preparation artifacts are much less likely when measuring protein-bound HNE because free HNE can be generated and sequestered very rapidly even during post-mortem lipid peroxidation. Such artifacts are well documented and have profound practical implications for seafood storage, durability, and flavor preservation (78, 79).

### Enrichment of the proteostasis machinery among HNE modified proteins

Ribosomal proteins and other proteins that are involved in proteostasis functions such as translation, protein stability, and protein turnover, are enriched in the set of partly HNE-modified proteins. This observation suggests that cellular proteostasis processes are specifically regulated by HNE protein adduction. Molecular chaperones are proteins central for proteostasis in all organisms (80) and among the HNE modified proteins identified in our study. HNE-adduction of molecular chaperones (HSP70 and HSP90 isoforms) in a human cancer cell line (RKO) has been suggested to represent a regulatory mechanism for activation of heat shock factor 1 (HSF1) and subsequent induction of cellular stress response genes that counteract oxidative stress and macromolecular damage (81). In rat livers treated with 10 - 100 μM HNE HSP72 adduction with HNE at the ATPase domain Cys267 increased significantly and inhibited HSP72 refolding (82). Rat liver cells also responded to HNE treatment by increased HNE adduction of HSP90 (51).

Consistent with a function of HNE adduction for regulating molecular chaperones, our study has identified HSP90-beta (chrUn.g68340.01: Lys270), HSP70 (chr10.g51517.01: His86, Lys123), HSP70.4 (chr5.g32332.01: Lys13, Arg43, His175, Arg178, Cys210, Arg314), HSP70.4b (chrUn.g32302.01: Lys86), HSC71 (chr4.g19408.01: Lys46, Cys578), mitochondrial HSP75 (chrUn.g16185.01: Cys30, Cys514), mitochondrial HSP75b (chrUn.g32390.02: Cys64, Cys888), BiP (chr5.g43996.01: Cys44, Arg52, Lys143, Lys184), and HSP60A (chr12.g64909.01: Cys488), as HNE adducted molecular chaperones in *B. schlosseri* (Supplementary table 1). We have previously shown that mRNA abundances for HSP90-beta and HSP70 isoforms increase during stress in *B. schlosseri* (83). Besides strengthening their role as molecular chaperones, elevation of HSP levels will also increase the capacity for sequestering HNE during stress.

In addition, HNE-adducted *B. schlosseri* molecular chaperones include a tubulin-specific chaperone (chr10.g68715.01: Arg954), T-complex protein 1 (TCP1) alpha (chrUn.g23678.01: Lys404), TCP1-gamma (chrUn.g10337.01: Lys140, Cys197, Lys203, Arg230, Lys240, Arg267, Arg343), TCP1-eta (chrUn.g30007.01: Lys319), TCP1-eta2 (chrUn.g51266.01, Lys266), TCP1-epsilon (chr5.g44307.01: Lys56, His82, Lys157), TCP1-theta (chr2.g69080.01: His8, Lys23, Arg104, Lys120, His207, Arg213, Arg220, Lys410, His460, Lys464), TCP1-zeta (chr2.g02303.01: Arg117, Arg317, Lys362, Cys369), TCP1-zeta2 (chrUn.g16096.01: Arg15, Lys60, Cys67, His169), and TCP1-zeta3 (chrUn.g29443.01: Arg117) (Supplementary table 1). Tubulin-associated chaperones assemble microtubules made of alpha/beta-tubulin heterodimers (84). The T-complex protein 1 (TCP1/TRiC) represents a molecular chaperonin having a ring- or barrel-shape structure similar to HSP60, which preferentially assists in the folding of cytoskeletal proteins (85, 86). The involvement of these molecular chaperons in regulating proteostasis and cytoskeletal functions is consistent with their enrichment among NHE-adducted *B. schlosseri* proteins.

We have identified 23 ribosomal proteins, 32 ubiqutin-proteolysis system proteins, and 10 additional proteins involved in translation control and protein synthesis that are HNE-adducted at 35, 45, and 17 amino acid residues, respectively (Supplemental table 1). These proteins contribute to proteostasis in addition to the molecular chaperones outlined above. Effects of HNE on ribosomal and translational control proteins have been reported previously for mammals. Eighteen translation-related proteins, including six ribosomal proteins, were significantly increased in abundance in response to HNE exposure of human colorectal carcinoma RKO cells (87). For adult rat cardiomyocytes it has been reported that exposure to 10 µM HNE for 1h stimulates protein synthesis by activation of the mammalian target of rapamycin (mTOR) and extracellular signal regulated kinases 1/2 (ERK1/2) signaling pathways and that HNE adduction of key proteins in these pathways is responsible for this effect (88). Our findings of overrepresentation of proteostasis proteins among NHE adducted *B. schlosseri* proteins in conjunction with the other studies discussed above indicate a critical role of HNE and oxidative stress in the regulation of protein synthesis, stability, and turnover.

### *B. schlosseri* proteins with population-specific HNE modification ratios

The *B. schlosseri* colonies reared for multiple blastogenic generations *ex situ* under controlled laboratory conditions, revealed an increased extent of HNE modification on partly modified amino acid residues relative to the two field populations. Comparing the Santa Barbara laboratory (C) versus Santa Barbara field (W) populations revealed nine significantly increased and no significantly decreased proteins. Comparing C versus the Des Moines field population (T) indicated four significantly increased and three significantly decreased proteins. Several of the proteins having significantly elevated HNE ratios in the laboratory population C over the two field populations (W and T) are involved in cytoskeletal organization and proteostasis. Overall, these results suggest that the extent of oxidative stress experienced by the laboratory reared population may be greater than that experienced by either field population. Elevated oxidative stress has also been observed in other animal models maintained in laboratory facilities, which can only partly reproduce the optimal conditions in the natural habitat (89). However, it should be noted that the laboratory population was kept at 18°C constant temperature year-round, which is optimal for metabolism. In contrast the field populations were approaching early fall and were perhaps reducing their oxidative metabolism. Moreover, in contrast to the laboratory population, the blastogenic cycle of different systems collected from the field populations was asynchronous.

The only HNE modification that was included in the DIA assay library and significantly different between the two field populations was on Arg49 of lamin-B1 (chr1.g14902.01). This modification was elevated in the Santa Barbara field population (W) over the Des Moines population (T). To the best of our knowledge, the present study is the first to report HNE modification of any lamin. Lamin-B1 is a nuclear protein pivotal for maintaining chromatin responsiveness to regulatory signals and environmental cues (90, 91). This multifunctional protein controls p53-dependent DNA repair, cell proliferation, senescence, gene expression, chromatin compactness, and structural organization of the nucleoskeleton (92, 93). Two previous reports mention HNE in the same context as lamin but not as a lamin PTM. A proteome study of V79-4 Chinese hamster lung cells found a 33% reduction of lamin-C after 24h treatment with 10 µM NHE (94). Another study found large concomitant increases of HNE and lamin in kidneys of Wister rats after ischemia-reperfusion stress (95). Moreover, several studies have shown that both lamin and HNE contribute to the regulation of apoptosis and other forms of programmed cell death (34, 35). Our results thus, suggest that lamin-B1 HNE modification contributes to the regulation gene expression, proliferation, senescence, and the blastogenic cycle in response to different environmental cues in the Santa Barbara (W) versus Des Moines (T) field populations.

### Conclusions

This is the first study demonstrating HNE modification of proteins in tunicates. The number of HNE modified proteins and the extent of HNE modification in the colonial tunicate *B. schlosseri* are both remarkably high. We present a mass spectrometry-based approach for reliably localizing HNE adducts to specific amino acid residues in tunicate proteins and for quantifying the ratio of HNE modification at these specific sites. The DIA assay library generated in this study and the HNE adducted proteins identified in laboratory and field populations of *B. schlosseri* empower comprehensive assessment of redox regulation and signaling in this species. Future follow-up investigations include biomonitoring other field populations exposed to stressful environments, studying how invasive populations conquer new and stressful environments, studies of laboratory populations kept under controlled conditions and exposed to specific environmental cues or during different stages of the blastogenic cycle, and studies directed at oxidative stress responses associated with transitions of tunicates and their cells from *in situ* to *ex situ* and *in vivo* to *in vitro* environments.

## DATA AVAILABILITY

The proteomics raw data, metadata, and quantitative data have been deposited in pertinent public databases (PanoramaPublic and ProteomeXchange). The PanoramaPublic permanent access link is https://panoramaweb.org/bosch02kl.url and the ProteomeXchange accession number is PXD050284. The dataset doi is: https://doi.org/10.6069/ek2v-gt71.

The reviewer login details for accessing these data are as follows (data will be made public upon manuscript acceptance):

Link: https://panoramaweb.org/bosch02kl.url

Login

Password: qQ4fb+gCSszx@8

## ACKNOWLEDGMENTS

We thank Drs. Ayelet Voskoboynik (Stanford Univ., Hopkins Marine Station) and Rick Grosberg (UC Davis) for their advice on B. schlosseri biology, culture, and genomic resources. We are also grateful for support from the Skyline and Panorama developer team (Univ. Washington, Seattle).

## AUTHOR CONTRIBUTIONS

D. Kültz performed proteome analyses, bioinformatics, data interpretation, funding, and wrote the manuscript. A.M. Gardell, A. De Tomaso, and G. Stoney organized sample collection, contributed to experimental design, funding, and edited the manuscript. B. Rinkevich and A. Qarri contributed to experimental design, data interpretation, funding, and edited the manuscript. J. Hamar contributed to sample preparation and edited the manuscript.

