## Supplementary figures and images for "Proteome-wide 4-hydroxy-2-nonenal signature of oxidative stress in the marine invasive tunicate *Botryllus schlosseri*"

### Supplementary Figure 1

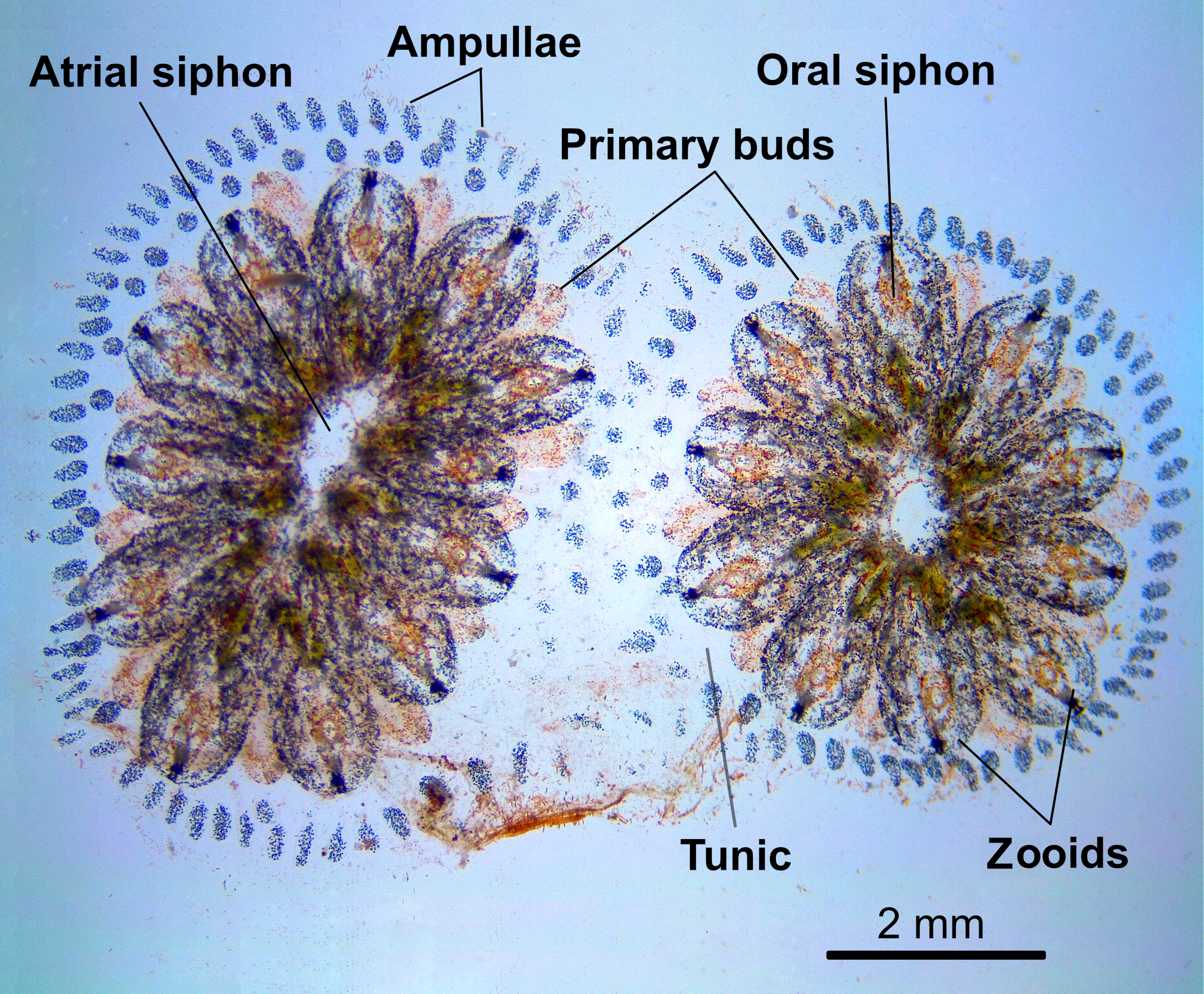
